## Supplementary Table S1 for "*MACREL*: antimicrobial peptide screening in genomes and metagenomes"

**Supplementary Table S1.** Features used in the MACREL classifiers.

| Descriptor | Definition | References | Variable Importance to classification of | |
|---|---|---|---|---|
| | | | AMPs[a] | Hemolytic peptides[a] |
| SA group 1 | Distribution parameter (CTDD) at first occurrence in the sequence using amino acids group 1 (A,L,F,C,G,I,V,W) clustered by solvent accessibility | Bhadra et al. (2018) | 26.2 | 12.0 |
| SA group 2 | Distribution parameter (CTDD) at first occurrence in the sequence using amino acids group 2 (R,K,Q,E,D,N) clustered by solvent accessibility | Bhadra et al. (2018) | 27.2 | 6.9 |
| SA group 3 | Distribution parameter (CTDD) at first occurrence in the sequence using amino acids group 3 (M,S,P,T,H,Y) clustered by solvent accessibility | Bhadra et al. (2018) | 100.0 | 8.4 |
| FET group 1 | Distribution parameter (CTDD) at first occurrence in the sequence using amino acids group 1 (I,L,V,W,A,G,T,M) clustered by the free energy to transfer from water to lipophilic phase | This study | 64.2 | 6.1 |
| FET group 2 | Distribution parameter (CTDD) at first occurrence in the sequence using amino acids group 2 (F,Y,S,Q,C,N) clustered by the free energy to transfer from water to lipophilic phase | This study | 16.9 | 6.1 |
| FET group 3 | Distribution parameter (CTDD) at first occurrence in the sequence using amino acids group 3 (P,H,K,E,D,R) clustered by the free energy to transfer from water to lipophilic phase | This study | 23.3 | 7.5 |
| Tiny res. % | Percent of residues of tiny amino acids (A + C + G + S + T) | Osorio et al. (2015) | 20.6 | 10.6 |
| Small res. % | Percent of residues of small amino acids (A + B + C + D + G + N + P + S + T + V) | Osorio et al. (2015) | 14.4 | 17.7 |
| Aliphatic res. % | Percent of residues of aliphatic amino acids (A + I + L + V) | Osorio et al. (2015) | 16.3 | 6.0 |
| Aromatic res. % | Percent of residues of aromatic amino acids (F + H + W + Y) | Osorio et al. (2015) | 13.1 | 7.6 |
| Nonpolar res. % | Percent of residues of nonpolar amino acids (A + C + F + G + I + L + M + P + V + W + Y) | Osorio et al. (2015) | 13.8 | 18.6 |
| Polar res. % | Percent of residues of polar amino acids (D + E + H + K + N + Q + R + S + T + Z) | Osorio et al. (2015) | 13.9 | 16.2 |
| Charged res. % | Percent of residues of charged amino acids (B + D + E + H + K + R + Z) | Osorio et al. (2015) | 12.6 | 9.6 |

| | | | | |
|---|---|---|---|---|
| Basic res. % | Percent of residues of basic amino acids (H + K + R) | Osorio et al. (2015) | 21.8 | 25.9 |
| Acidic res. % | Percent of residues of acidic amino acids (B + D + E + Z); | Osorio et al. (2015) | 31.9 | 79.0 |
| Peptide charge | Peptide charge at pH 7.0 using "EMBOSS" pKscale | EMBOSS | 34.5 | 88.4 |
| Isoelectric point | Peptide isoelectric point using "EMBOSS" pKscale | Bjellqvist et al. (1994) | 29.9 | 42.8 |
| Aliphatic index | Relative volume occupied by aliphatic side chains (A, V, I, and L) in the peptide chain | Ikai (1980) | 17.5 | 7.9 |
| Instability index | Peptide's stability of a protein based on its dipeptide composition, it is used to determine whether the peptide will be stable in a test tube. | Guruprasad et al. (1990) | 20.8 | 10.3 |
| Boman index | Sum of the solubility values for all residues in a sequence, it might give an overall estimate of the potential of a peptide to bind to membranes or other proteins as receptors, to normalize it is divided by the number of residues | Boman (2003) | 18.4 | 21.0 |
| Hydrophobicity | Peptide's hydrophobicity using "KyteDoolittle" scale | Kyte and Doolitle (1982) | 15.7 | 11.0 |
| H-moment | Quantitative measure of the amphiphilicity perpendicular to the axis of any periodic peptide structure, such as the alpha-helix or beta-sheet using angle of 100º and a window of 11 residues | Eisenberg et al. (1984) | 24.0 | 21.8 |

[a] Variable importance is a measure that refers to the frequency a given variable is used by the trees in the random forest
