## Supplementary Table S2 for "*MACREL*: antimicrobial peptide screening in genomes and metagenomes"

**Supplementary Table S2.** Homology and AMPs proportion in the training set affect different classifiers.

| Model/Method | NAMP/AMP | Accuracy | Specificity | Sensitivity | Precision | F-Score | MCC |
|---|---|---|---|---|---|---|---|
| Homology Classification | 1 | 0.222 | 0.136 | 0.308 | 0.263 | 0.284 | -0.564 |
| Homology Classification | 5 | 0.318 | 0.346 | 0.290 | 0.307 | 0.298 | -0.365 |
| Homology Classification | 10 | 0.352 | 0.416 | 0.288 | 0.330 | 0.308 | -0.298 |
| Homology Classification | 20 | 0.384 | 0.498 | 0.270 | 0.350 | 0.305 | -0.238 |
| Homology Classification | 30 | 0.421 | 0.580 | 0.262 | 0.384 | 0.312 | -0.167 |
| Homology Classification | 40 | 0.432 | 0.600 | 0.264 | 0.398 | 0.317 | -0.144 |
| Homology Classification | 50 | 0.437 | 0.622 | 0.252 | 0.400 | 0.309 | -0.136 |
| Macrel Re-trained | 1 | 0.926 | 0.952 | 0.900 | 0.949 | 0.924 | 0.853 |
| Macrel Re-trained | 5 | 0.893 | 0.986 | 0.800 | 0.983 | 0.882 | 0.800 |
| Macrel Re-trained | 10 | 0.849 | 0.996 | 0.702 | 0.994 | 0.823 | 0.730 |
| Macrel Re-trained | 20 | 0.804 | 0.996 | 0.612 | 0.994 | 0.757 | 0.658 |
| Macrel Re-trained | 30 | 0.763 | 1.000 | 0.526 | 1.000 | 0.689 | 0.597 |
| Macrel Re-trained | 40 | 0.734 | 1.000 | 0.468 | 1.000 | 0.638 | 0.553 |
| Macrel Re-trained | 50 | 0.718 | 1.000 | 0.436 | 1.000 | 0.607 | 0.528 |
| AMP Scanner Re-trained | 1 | 0.908 | 0.926 | 0.890 | 0.923 | 0.906 | 0.817 |
| AMP Scanner Re-trained | 5 | 0.923 | 0.956 | 0.890 | 0.953 | 0.920 | 0.848 |
| AMP Scanner Re-trained | 10 | 0.894 | 0.984 | 0.804 | 0.980 | 0.884 | 0.801 |
| AMP Scanner Re-trained | 20 | 0.889 | 0.994 | 0.784 | 0.992 | 0.876 | 0.796 |
| AMP Scanner Re-trained | 30 | 0.845 | 0.996 | 0.694 | 0.994 | 0.817 | 0.724 |
| AMP Scanner Re-trained | 40 | 0.856 | 0.990 | 0.722 | 0.986 | 0.834 | 0.739 |
| AMP Scanner Re-trained | 50 | 0.850 | 0.992 | 0.708 | 0.989 | 0.825 | 0.730 |
| iAMP-2L Re-trained | 1 | 0.924 | 0.920 | 0.928 | 0.921 | 0.924 | 0.848 |
| iAMP-2L Re-trained | 5 | 0.908 | 0.980 | 0.836 | 0.977 | 0.901 | 0.825 |
| iAMP-2L Re-trained | 10 | 0.894 | 0.990 | 0.798 | 0.988 | 0.883 | 0.803 |
| iAMP-2L Re-trained | 20 | 0.871 | 0.996 | 0.746 | 0.995 | 0.853 | 0.766 |
| iAMP-2L Re-trained | 30 | 0.849 | 1.000 | 0.698 | 1.000 | 0.822 | 0.732 |
| iAMP-2L Re-trained | 40 | 0.824 | 1.000 | 0.648 | 1.000 | 0.786 | 0.692 |

| Model/Method | NAMP/AMP | Accuracy | Specificity | Sensitivity | Precision | F-Score | MCC |
|---|---|---|---|---|---|---|---|
| iAMP-2L Re-trained | 50 | 0.811 | 1.000 | 0.622 | 1.000 | 0.767 | 0.672 |
