## Supplementary Table S3 for "*MACREL*: antimicrobial peptide screening in genomes and metagenomes"

**Supplementary Table S3.** MACREL analysis using real metagenomes from human guts (Heinz et al., 2016). The SRR code shows the access to the metagenome in SRA database. The number of predicted smORFs, AMPs and spurious predictions after AMP classification are shown. Total length, GC(%) and N50 were measured after filtering contigs by minimum length of 1kpb.

| SRR | # contigs | Total length (Mbp) ** | GC (%) | N50 | smORFs | AMPs | Spurious |
|---|---|---|---|---|---|---|---|
| 3313034 | 22286 | 77.89 | 43.34 | 5594 | 12727 | 49 | 6 |
| 3313035 | 37040 | 125.18 | 41.41 | 5023 | 20318 | 83 | 4 |
| 3313036 | 37786 | 141.80 | 46.13 | 6588 | 23220 | 94 | 1 |
| 3313037 | 19931 | 85.93 | 43.53 | 10679 | 13326 | 58 | 1 |
| 3313038 | 43246 | 154.67 | 44 | 5913 | 23924 | 83 | 4 |
| 3313039 | 30019 | 103.59 | 43.89 | 5700 | 16743 | 60 | 1 |
| 3313040 | 26025 | 110.99 | 43.59 | 9581 | 17010 | 57 | 6 |
| 3313041 | 29712 | 111.51 | 43.62 | 6675 | 17054 | 66 | 5 |
| 3313042 | 29339 | 112.75 | 44.18 | 6739 | 16458 | 54 | 5 |
| 3313043 | 24661 | 89.07 | 45.93 | 5673 | 14315 | 41 | 0 |
| 3313044 | 33010 | 114.55 | 45.34 | 5267 | 18614 | 66 | 5 |
| 3313045 | 30004 | 109.24 | 44.4 | 5846 | 17596 | 49 | 4 |
| 3313046 | 41867 | 135.27 | 42.78 | 5098 | 23634 | 80 | 7 |
| 3313047 | 25340 | 91.32 | 45.94 | 5761 | 14496 | 59 | 7 |
| 3313048 | 31340 | 115.55 | 44.52 | 5781 | 17777 | 61 | 5 |
| 3313049 | 29247 | 108.09 | 45.23 | 6089 | 16198 | 69 | 7 |
| 3313050 | 43878 | 151.64 | 45.65 | 5281 | 23388 | 80 | 10 |
| 3313051 | 33168 | 120.45 | 46.17 | 6549 | 18449 | 61 | 9 |
| 3313052 | 39579 | 154.58 | 45.97 | 7331 | 22457 | 83 | 5 |
| 3313053 | 5613 | 28.49 | 42.94 | 12389 | 3879 | 18 | 1 |
| 3313054 | 18504 | 108.82 | 43.88 | 19575 | 14517 | 70 | 3 |
| 3313055 | 41163 | 146.14 | 43.98 | 5813 | 22789 | 81 | 1 |
| 3313056 | 43301 | 151.58 | 44.1 | 5573 | 27480 | 83 | 6 |
| 3313057 | 34906 | 133.87 | 43.79 | 7408 | 19365 | 78 | 7 |
| 3313058 | 25605 | 121.19 | 44.89 | 9415 | 16884 | 63 | 6 |
| 3313059 | 36287 | 137.11 | 44.4 | 6256 | 19761 | 81 | 8 |
| 3313060 | 34833 | 132.45 | 45.05 | 6348 | 18776 | 72 | 4 |
| 3313061 | 26900 | 128.26 | 43.97 | 14264 | 17414 | 72 | 2 |
| 3313062 | 35491 | 137.62 | 44.38 | 7873 | 20064 | 82 | 7 |
| 3313063 | 36324 | 153.81 | 44.42 | 8284 | 22078 | 97 | 9 |
| 3313064 | 17535 | 77.29 | 43.48 | 11892 | 11207 | 38 | 4 |
| 3313065 | 20943 | 114.53 | 45.27 | 16631 | 14498 | 73 | 7 |
| 3313066 | 16373 | 65.69 | 46.68 | 7679 | 9723 | 51 | 2 |
| 3313067 | 23158 | 92.79 | 43.7 | 7593 | 13754 | 56 | 3 |
| 3313068 | 28706 | 135.19 | 43.59 | 12173 | 19388 | 82 | 4 |
| 3313069 | 24284 | 87.50 | 44.81 | 6074 | 13601 | 51 | 4 |
| 3313070 | 31368 | 127.33 | 46.65 | 7878 | 18706 | 96 | 4 |
| 3313071 | 22612 | 66.16 | 44.9 | 3825 | 11526 | 41 | 4 |
| 3313072 | 33597 | 124.53 | 45.37 | 6280 | 18970 | 86 | 8 |
| 3313073 | 20705 | 79.42 | 44.13 | 6722 | 11909 | 51 | 3 |
| 3313074 | 29127 | 107.98 | 44.21 | 6195 | 16508 | 63 | 12 |

| SRR | # contigs | Total length (Mbp) ** | GC (%) | N50 | smORFs | AMPs | Spurious |
|---|---|---|---|---|---|---|---|
| 3313075 | 20946 | 74.54 | 42.99 | 5736 | 11763 | 51 | 6 |
| 3313076 | 17748 | 84.78 | 45.87 | 11344 | 11908 | 57 | 5 |
| 3313077 | 23590 | 117.26 | 44.91 | 12372 | 16011 | 73 | 5 |
| 3313078 | 20259 | 75.94 | 43.03 | 6449 | 11872 | 51 | 6 |
| 3313079 | 35547 | 141.09 | 42.28 | 7274 | 20972 | 81 | 12 |
| 3313080 | 24686 | 83.04 | 45.44 | 5599 | 12230 | 58 | 4 |
| 3313081 | 23832 | 87.43 | 42.16 | 6326 | 14179 | 74 | 1 |
| 3313082 | 36846 | 132.37 | 44.9 | 5692 | 21264 | 94 | 3 |
| 3313090 | 30188 | 113.75 | 44.14 | 7091 | 17117 | 58 | 2 |
| 3313102 | 37144 | 127.73 | 43.86 | 5411 | 20320 | 65 | 2 |
| 3313113 | 31894 | 119.28 | 45.2 | 6087 | 17846 | 68 | 4 |
| 3313123 | 38138 | 128.81 | 43.95 | 5060 | 22346 | 87 | 2 |
| 8114009 | 28753 | 78.93 | 40.61 | 3334 | 13302 | 39 | 3 |
| 8114010 | 44483 | 123.33 | 41.77 | 3368 | 21064 | 66 | 11 |
| 8114011 | 3336 | 9.07 | 47.43 | 3329 | 1490 | 2 | 0 |
| 8114012 | 4889 | 11.67 | 52.2 | 2515 | 1813 | 2 | 0 |
| 8114013 | 28567 | 83.80 | 41.31 | 3821 | 15044 | 37 | 1 |
| 8114014 | 26822 | 73.51 | 39.78 | 3317 | 12976 | 33 | 1 |
| 8114015 | 3697 | 6.12 | 57.29 | 1648 | 1146 | 0 | 0 |
| 8114016 | 13063 | 34.60 | 39.82 | 3180 | 6441 | 18 | 2 |
| 8114017 | 13910 | 42.82 | 46.6 | 4189 | 6673 | 14 | 0 |
| 8114018 | 959 | 1.69 | 38.97 | 1756 | 401 | 2 | 0 |
| 8114019 | 9441 | 27.45 | 56.14 | 3735 | 3921 | 9 | 0 |
| 8114020 | 2672 | 5.40 | 39.06 | 2096 | 1133 | 5 | 0 |
| 8114021 | 12636 | 36.66 | 42.57 | 3563 | 5879 | 17 | 2 |
| 8114022 | 4642 | 11.12 | 60.45 | 2799 | 1622 | 5 | 0 |
| 8114023 | 2370 | 4.40 | 43.71 | 1870 | 1027 | 1 | 0 |
| 8114024 | 10209 | 21.33 | 49.77 | 2149 | 4406 | 13 | 4 |
| 8114025 | 59573 | 258.87 | 46.95 | 9853 | 38004 | 165 | 9 |
| 8114026 | 48294 | 201.95 | 46.72 | 9166 | 29731 | 122 | 10 |
| 8114027 | 50975 | 212.54 | 46.64 | 8959 | 31906 | 137 | 7 |
| 8114028 | 31965 | 110.55 | 44.56 | 5381 | 16084 | 62 | 4 |
| 8114029 | 3115 | 5.66 | 60.2 | 1841 | 950 | 0 | 0 |
| 8114030 | 21466 | 60.49 | 52.63 | 3598 | 9146 | 37 | 6 |
| 8114031 | 833 | 1.23 | 62.24 | 1449 | 243 | 4 | 0 |
| 8114032 | 1191 | 1.70 | 46.98 | 1361 | 403 | 0 | 0 |
| 8114033 | 22109 | 59.50 | 53.21 | 3176 | 9104 | 34 | 8 |
| 8114034 | 23468 | 81.28 | 49.18 | 5146 | 10644 | 35 | 4 |
| 8114035 | 14533 | 42.43 | 45.99 | 3628 | 6901 | 31 | 7 |
| 8114036 | 10762 | 29.40 | 48.96 | 3236 | 4733 | 11 | 1 |
| 8114037 | 6987 | 22.22 | 44.11 | 4575 | 3557 | 7 | 0 |
| 8114038 | 21 | 0.03 | 45.05 | 1700 | 7 | 0 | 0 |
| 8114039 | 19253 | 51.14 | 45.15 | 3085 | 8068 | 14 | 1 |
| 8114040 | 10067 | 20.89 | 41.94 | 2248 | 3647 | 7 | 1 |
| 8114041 | 9863 | 22.85 | 39.65 | 2464 | 4272 | 14 | 2 |
| 8114042 | 4945 | 10.93 | 37.29 | 2404 | 2192 | 6 | 0 |
| 8114043 | 6344 | 15.55 | 65.45 | 2798 | 2243 | 3 | 1 |
| 8114044 | 6050 | 12.71 | 40.15 | 2245 | 2415 | 5 | 1 |
| 8114045 | 11208 | 33.95 | 48.02 | 4018 | 4630 | 16 | 1 |

| SRR | # contigs | Total length (Mbp) ** | GC (%) | N50 | smORFs | AMPs | Spurious |
|---|---|---|---|---|---|---|---|
| 8114046 | 702 | 1.44 | 56.15 | 1881 | 332 | 1 | 0 |
| 8114047 | 7056 | 18.36 | 60.26 | 3009 | 2925 | 5 | 0 |
| 8114048 | 8762 | 19.60 | 39.45 | 2289 | 3932 | 6 | 1 |
| 8114049 | 9327 | 25.87 | 42.21 | 3413 | 4417 | 13 | 3 |
| 8114050 | 626 | 1.48 | 40.68 | 2618 | 507 | 0 | 0 |
| 8114051 | 23777 | 60.76 | 41.65 | 2900 | 10113 | 33 | 3 |
| 8114052 | 39553 | 101.95 | 38.78 | 3015 | 19009 | 54 | 3 |
| 8114053 | 17084 | 60.55 | 46.2 | 5417 | 8943 | 32 | 2 |
| 8114054 | 15769 | 54.37 | 43.5 | 5145 | 8420 | 21 | 0 |
| 8114055 | 14297 | 46.73 | 52.69 | 4905 | 6391 | 17 | 1 |
| 8114056 | 21120 | 66.16 | 45.04 | 4298 | 11565 | 33 | 2 |
| 8114057 | 7262 | 16.66 | 40.78 | 2512 | 2951 | 10 | 1 |
| 8114058 | 12264 | 49.86 | 43.45 | 8673 | 7336 | 32 | 2 |
| 8114059 | 1777 | 4.40 | 39.4 | 2839 | 795 | 9 | 0 |
| 8114060 | 31680 | 90.00 | 41.81 | 3477 | 13786 | 44 | 3 |
| 8114061 | 293 | 0.48 | 39.47 | 1452 | 138 | 1 | 0 |
| 8114062 | 19259 | 45.93 | 40.8 | 2542 | 7577 | 27 | 2 |
| 8114063 | 30350 | 82.29 | 42.33 | 3223 | 13351 | 50 | 3 |
| 8114064 | 25997 | 75.59 | 47.66 | 3702 | 10767 | 30 | 5 |
| 8114065 | 6924 | 17.15 | 47.83 | 2862 | 2683 | 7 | 1 |
| 8114066 | 3989 | 7.61 | 39.92 | 1894 | 1639 | 3 | 1 |
| 8114067 | 20452 | 54.78 | 58.91 | 3185 | 8377 | 31 | 3 |
| 8114068 | 5838 | 13.29 | 46.24 | 2378 | 2399 | 5 | 0 |
| 8114069 | 9633 | 19.69 | 39.63 | 1970 | 4055 | 10 | 2 |
| 8114070 | 21202 | 56.25 | 41.38 | 3133 | 9448 | 38 | 5 |
| 8114071 | 17145 | 37.88 | 40.88 | 2222 | 6713 | 22 | 1 |
| 8114072 | 14880 | 47.14 | 48.63 | 4323 | 7140 | 21 | 6 |
| 8114073 | 21404 | 60.52 | 44.59 | 3648 | 9291 | 37 | 2 |
| 8114074 | 346 | 0.50 | 40.86 | 1398 | 135 | 1 | 0 |
| 8114075 | 8124 | 20.05 | 51.39 | 2779 | 3240 | 13 | 1 |
| 8114076 | 27221 | 78.97 | 46.33 | 3630 | 12340 | 45 | 4 |
| 8114077 | 23941 | 66.94 | 39.83 | 3395 | 11466 | 31 | 4 |
| 8114078 | 26777 | 65.05 | 40.24 | 2688 | 12237 | 32 | 5 |
| 8114079 | 8914 | 17.14 | 37.12 | 1864 | 3869 | 8 | 0 |
| 8114080 | 7746 | 22.77 | 58.61 | 3781 | 3065 | 8 | 1 |
| 8114081 | 4202 | 10.82 | 66.46 | 3221 | 1476 | 2 | 1 |
| 8114082 | 6975 | 19.46 | 48.39 | 3268 | 2964 | 10 | 1 |
| 8114083 | 7601 | 25.60 | 48.77 | 4603 | 3651 | 6 | 0 |
| 8114084 | 20045 | 70.20 | 41.63 | 5194 | 10595 | 29 | 1 |
| 8114085 | 10322 | 27.51 | 46.93 | 3112 | 4913 | 13 | 3 |
| 8114086 | 12904 | 40.21 | 41.49 | 4314 | 6565 | 22 | 4 |
| 8114087 | 16569 | 51.59 | 50.04 | 4179 | 7229 | 21 | 0 |
| 8114088 | 17896 | 47.28 | 43.74 | 3122 | 7826 | 20 | 1 |
| 8114089 | 7424 | 15.58 | 42.88 | 2201 | 3228 | 5 | 1 |
| 8114090 | 5273 | 13.08 | 53.29 | 2873 | 1931 | 8 | 2 |
| 8114091 | 16960 | 46.21 | 43.04 | 3338 | 7122 | 16 | 1 |
| 8114092 | 19841 | 48.45 | 37.55 | 2750 | 9200 | 29 | 2 |
| 8114093 | 17665 | 38.63 | 40.15 | 2263 | 7569 | 17 | 1 |
| 8114094 | 25101 | 66.52 | 40.13 | 3109 | 11907 | 46 | 1 |

| SRR | # contigs | Total length (Mbp) ** | GC (%) | N50 | smORFs | AMPs | Spurious |
|---|---|---|---|---|---|---|---|
| 8114095 | 15459 | 36.97 | 45.69 | 2593 | 6327 | 13 | 1 |
| 8114096 | 22138 | 78.59 | 41.36 | 6052 | 12475 | 49 | 5 |
| 8114097 | 17219 | 46.36 | 50.3 | 3139 | 6706 | 25 | 3 |
| 8114098 | 30307 | 156.19 | 46.35 | 15594 | 20133 | 100 | 13 |
| 8114099 | 36067 | 144.43 | 49.71 | 7428 | 17831 | 79 | 7 |
| 8114100 | 42306 | 181.81 | 47.28 | 9223 | 25805 | 134 | 7 |
| 8114101 | 30373 | 144.67 | 46.95 | 11060 | 20398 | 93 | 4 |
| 8114102 | 47499 | 196.94 | 46.62 | 7605 | 27274 | 121 | 1 |
| 8114103 | 33665 | 178.91 | 47.17 | 15294 | 23227 | 104 | 10 |
| 8114104 | 49645 | 185.82 | 47.99 | 6448 | 27172 | 121 | 9 |
| 8114105 | 49906 | 199.41 | 45.27 | 7382 | 28963 | 130 | 7 |
| 8114106 | 35185 | 139.64 | 45.33 | 7756 | 20467 | 87 | 8 |
| 8114107 | 46543 | 220.48 | 47.31 | 11709 | 28492 | 127 | 11 |
| 8114108 | 20196 | 99.58 | 48.07 | 19068 | 12774 | 57 | 3 |
| 8114109 | 36286 | 174.10 | 45.74 | 11630 | 23027 | 107 | 6 |
| 8114110 | 27166 | 120.79 | 46.82 | 9321 | 17293 | 67 | 3 |
| 8114111 | 32818 | 133.63 | 46.61 | 7635 | 20079 | 83 | 7 |
| 8114112 | 34043 | 141.58 | 45.35 | 8039 | 21114 | 99 | 12 |
| 8114113 | 44638 | 180.33 | 46.83 | 7040 | 25662 | 110 | 9 |
| 8114114 | 40288 | 184.70 | 47.67 | 11374 | 26002 | 132 | 11 |
| 8114115 | 35747 | 167.62 | 46.6 | 10865 | 23162 | 110 | 4 |
| 8114116 | 23490 | 101.91 | 47.22 | 8459 | 13894 | 59 | 3 |
| 8114117 | 32080 | 129.05 | 44.92 | 7949 | 20838 | 96 | 7 |
| 8114118 | 41909 | 156.06 | 47.3 | 6152 | 22116 | 104 | 9 |
| 8114119 | 43531 | 133.91 | 51.95 | 4144 | 18850 | 81 | 8 |
| 8114120 | 20955 | 86.61 | 46.17 | 8922 | 12867 | 63 | 3 |
| 8114121 | 30344 | 121.21 | 46.46 | 6738 | 17466 | 80 | 2 |
| 8114122 | 18776 | 104.19 | 46.12 | 20637 | 13987 | 55 | 9 |
| 8114123 | 11866 | 60.85 | 45.53 | 11302 | 7785 | 37 | 6 |
| 8114124 | 26423 | 113.65 | 47.14 | 8765 | 15261 | 66 | 13 |
| 8114125 | 24806 | 123.27 | 47.32 | 15989 | 16459 | 68 | 9 |
| 8114126 | 24388 | 80.45 | 50.47 | 4971 | 12187 | 61 | 8 |
| 8114127 | 22766 | 128.24 | 47.65 | 14761 | 16278 | 71 | 6 |
| 8114128 | 46594 | 182.97 | 48.89 | 7280 | 24591 | 106 | 11 |
| 8114129 | 39040 | 160.37 | 46.97 | 7639 | 20930 | 115 | 13 |
| 8114130 | 51552 | 194.59 | 47.74 | 6998 | 29983 | 124 | 6 |
| 8114131 | 41099 | 169.02 | 45.85 | 8300 | 23586 | 93 | 2 |
| 8114132 | 41762 | 191.28 | 46.15 | 10656 | 25900 | 121 | 10 |
| 8114133 | 48579 | 198.94 | 47.11 | 7951 | 28175 | 131 | 10 |
| 8114134 | 37515 | 173.22 | 48.23 | 10391 | 23539 | 107 | 9 |
| 8114135 | 39256 | 159.80 | 47.84 | 7507 | 21729 | 83 | 6 |
| 8114136 | 36389 | 176.72 | 47.2 | 12266 | 23943 | 104 | 8 |
| 8114137 | 28765 | 127.71 | 46.13 | 9702 | 17527 | 81 | 9 |

**Total Length – sum of length of contigs longer than 1 kbp given in Mbp ($10^6$ bp).
