## Supplementary Table S4 for "*MACREL*: antimicrobial peptide screening in genomes and metagenomes"

**Supplementary Table S4.** Co-prediction of MACREL results from real metagenomes with other methods. The AMPs predicted by MACREL were sorted into co-predicted AMPs (Agree column) and those predicted as NAMP by other methods (Disagree column). Co-prediction level was calculated as the percent of agreement with MACREL and the highest co-prediction level is bold.

| Method | Algorithm[b] | # Agree | # Disagree | Co-prediction |
|---|---|---|---|---|
| AMP Scanner v.2 | NN | 2068 | 1866 | 52.57 |
| CAMPR3 | SVM | 2334 | 1600 | 59.33 |
| CAMPR3 | RF | 2541 | 1393 | 64.59 |
| CAMPR3 | NN | 2538 | 1396 | 64.51 |
| CAMPR3 | DA | 2451 | 1483 | 62.30 |
| iAMPpred | SVM | 2777 | 1157 | **70.59** |
| AMAP | SVM | 2056 | 1878 | 52.26 |
| iAMP-2L | FKNN | 1350 | 2584 | 34.32 |
| Overall | >= 4 methods | 2549 | 1385 | 64.79 |
| Min. | >= 1 method | 3604 | 330 | **91.61** |

[a] Overall results show the number of peptides predicted as AMP by at least the half (4) of the methods simultaneously;

[b] Algorithm used by the method: SVM – Supported vector machines. FKNN – K-nearest neighborhood. NN – Neural network. DA – Discriminant analysis. RF – Random forest.
