## Supplementary Table S5 for "*MACREL*: antimicrobial peptide screening in genomes and metagenomes"

**Supplementary Table S5.** Datasets used in training and testing of classifiers implemented in Macrel.

| Dataset | Positives | Negatives | Neg. : Pos. | Positives Length distribution (%) | | | | | Negatives Length distribution (%) | | | | | Overlap[1] | Source | Experiment |
|---|---|---|---|---|---|---|---|---|---|---|---|---|---|---|---|---|
| | | | | 0-25 | 25-50 | 50-75 | 75-100 | + | 0-25 | 25-50 | 50-75 | 75-100 | + | | | |
| Trainingset_AMPs_1_1 | 1197 | 1197 | 1 | | | | | | 1,5 | 3,0 | 5,8 | 9,9 | 79,7 | | Bhadra et al. (2018) | |
| Trainingset_AMPs_1_5 | 1197 | 6000 | 5 | | | | | | 1,2 | 2,1 | 5,4 | 10,0 | 81,3 | | Bhadra et al. (2018) | |
| Trainingset_AMPs_1_10 | 1197 | 12000 | 10 | | | | | | 1,3 | 2,3 | 5,5 | 9,6 | 81,3 | | Bhadra et al. (2018) | |
| Trainingset_AMPs_1_20 | 1197 | 24000 | 20 | 36,3 | 39,1 | 10,9 | 4,8 | 8,9 | 1,2 | 2,1 | 5,8 | 9,9 | 80,9 | 1_5,1_10,1_20,1_30,1_40,1_50 | Bhadra et al. (2018) | |
| Trainingset_AMPs_1_30 | 1197 | 36000 | 30 | | | | | | 1,3 | 2,2 | 5,7 | 9,9 | 80,8 | | Bhadra et al. (2018) | Homology |
| Trainingset_AMPs_1_40 | 1197 | 48000 | 40 | | | | | | 1,3 | 2,1 | 5,6 | 10 | 80,9 | | Bhadra et al. (2018) | |
| Trainingset_AMPs_1_50 | 1197 | 60000 | 50 | | | | | | 1,4 | 2,2 | 5,6 | 10,0 | 80,8 | | Bhadra et al. (2018) | |
| Testingset_AMPs_1_1 | 500 | 500 | 1 | 34,4 | 41,6 | 8,4 | 5,6 | 10 | 0,6 | 2 | 6,2 | 11,4 | 79,8 | None | Bhadra et al. (2018) | |
| AMP.test | 920 | 920 | 1 | 44,7 | 52 | 3,3 | - | - | 10,6 | 3,7 | 19,4 | 66,3 | - | AMP.train | Xiao et al. (2013) | |
| AMP.train_bench | 1476 | 2405 | 2 | 48,3 | 42,9 | 5,4 | 3,3 | 0,1 | 10,2 | 11,2 | 29 | 49,6 | - | None | Xiao et al. (2013) | |
| AMP.train | 3268 | 165138 | 50 | 37,5 | 40,8 | 8,7 | 3,7 | 9,3 | 1,5 | 2,5 | 5,6 | 10,1 | 80,3 | AMP.test | Bhadra et al. (2018) | Macrel model |
| Hemo.train | 442 | 442 | 1 | 76,0 | 23,3 | 0,4 | 0,3 | - | 74,7 | 22,4 | 2,2 | 0,7 | - | None | Chaudhary et al. (2016) | |
| Hemo.test | 110 | 110 | 1 | 72,7 | 27,3 | - | - | - | 78,2 | 19,1 | 1,8 | 0,9 | - | None | Chaudhary et al. (2016) | |

[1] Overlap in terms of homologous peptides
