## Supplementary Figure S1 for "*MACREL*: antimicrobial peptide screening in genomes and metagenomes"

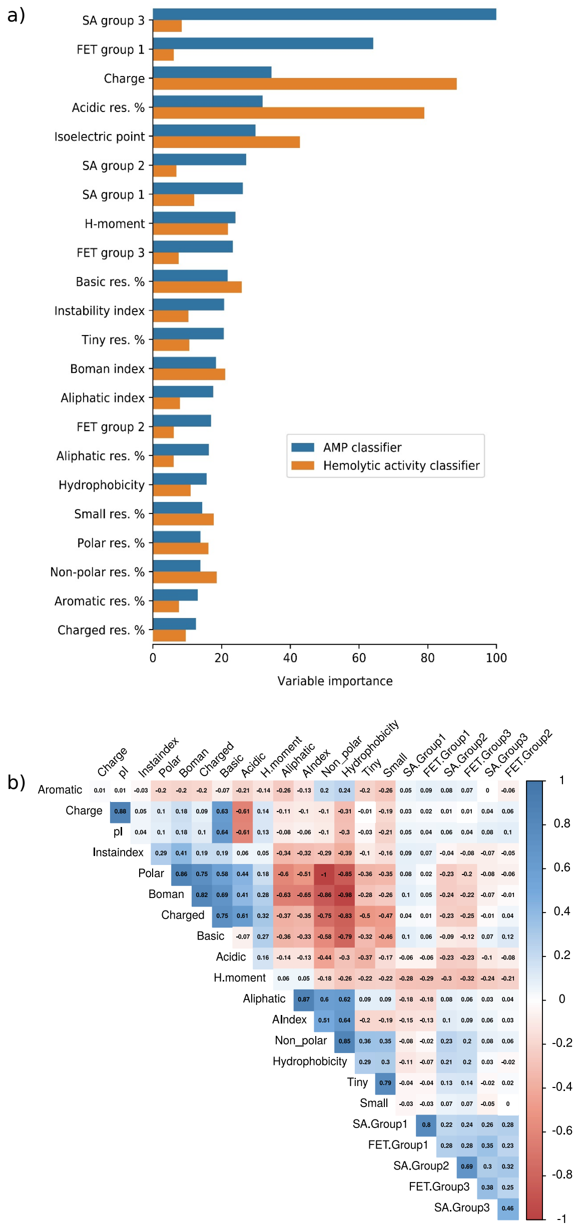


**Supplementary Figure S1. All 22 features are important in classification.** (a) Variable importance is measured as the percentage of times each variable is selected in the pruned models (MACREL classifiers of AMPs and hemolytic peptides). The solvent accessibility and free energy to transfer from water to lipophilic phase residues distribution at first position using 3 amino acid groups were summarized as SA and FET, respectively. (b) A correlogram showing the Pearson's correlation coefficient of the different variables used in our classifiers is shown.

Statistical significant ($p < 0.05$) correlations were shown, white boxes denote insignificant correlations.
