## Supplementary Figure S2 for "*MACREL*: antimicrobial peptide screening in genomes and metagenomes"

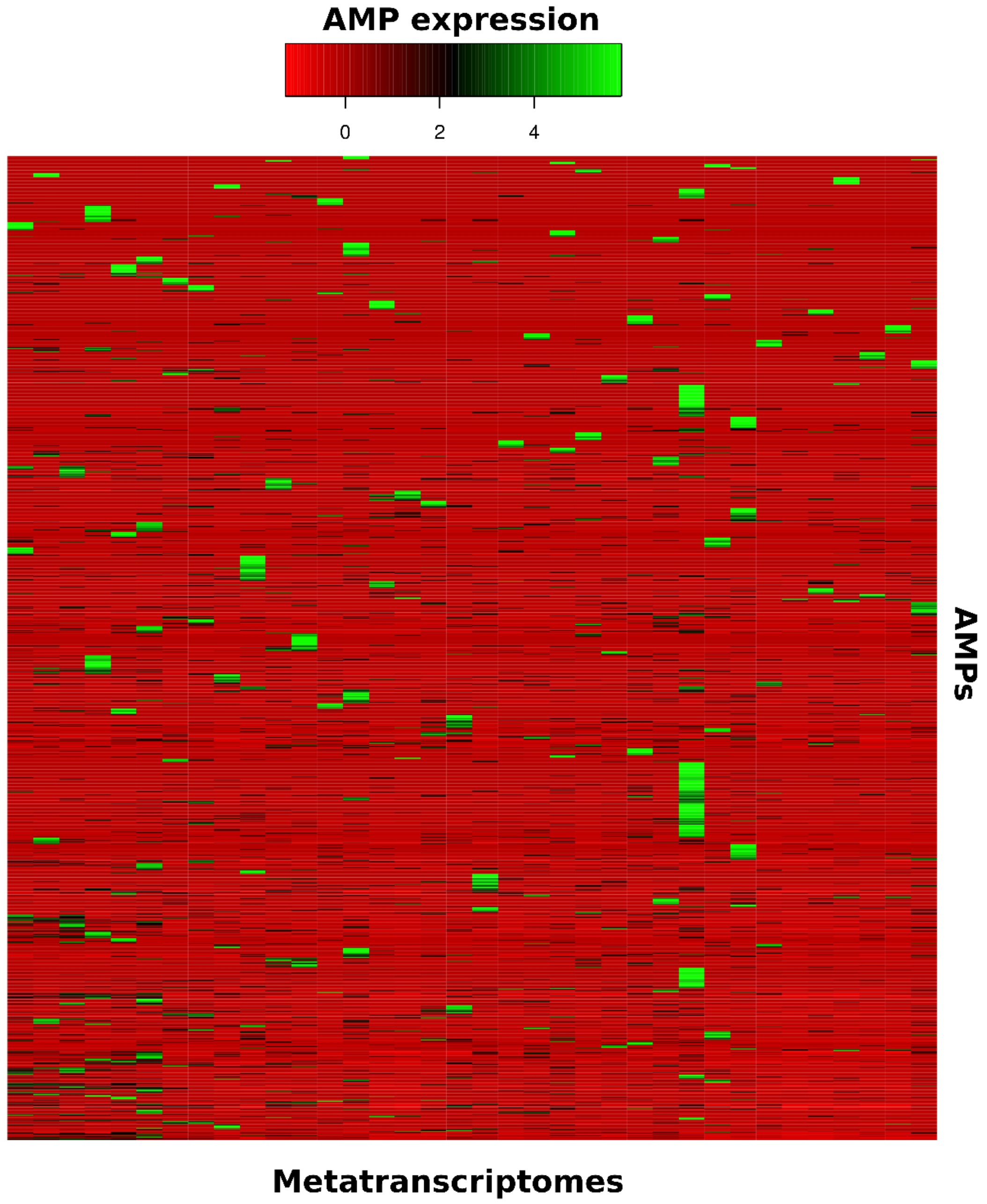


**Supplementary Figure S2. Gene expression of AMPs detected by MACREL in human gut metatranscriptomes.** The heatmap shows the AMPs with at least one of the expression values different from zero, and normalized by metatranscriptome using Z-score.
